# Co-occurrence of essential gene dispensability and bypass suppressor mutations across species

**DOI:** 10.1101/2022.11.11.516117

**Authors:** Carles Pons, Jolanda van Leeuwen

## Abstract

Genes have been historically classified as either essential or non-essential based on their requirement for viability. However, some genes are essential in some genetic backgrounds but non-essential in others, thus challenging the binary classification of gene essentiality. Such dispensable essential genes represent a valuable model for understanding the incomplete penetrance of loss-of-function mutations that is often observed in natural populations. Here, we compiled data from multiple studies on essential gene dispensability in *Saccharomyces cerevisiae* to comprehensively characterize these genes. In analyses spanning different evolutionary time-scales, ranging from *S. cerevisiae* strains to human cell lines, dispensable essential genes exhibited distinct phylogenetic properties compared to other essential and non-essential genes. Integration of interactions with suppressor genes that can bypass the gene essentiality revealed the high functional modularity of the bypass suppression network. Furthermore, dispensable essential and bypass suppressor gene pairs reflected simultaneous changes in the mutational landscape of *S. cerevisiae* strains. Importantly, species in which dispensable essential genes were non-essential tended to carry bypass suppressor mutations in their genomes. Overall, our study offers a comprehensive view of dispensable essential genes and illustrates how their interactions with bypass suppressor genes reflect evolutionary outcomes.

## INTRODUCTION

Identification of the genes required for viability is key for both fundamental and applied biological research. Essential genes constrain genome evolution (Jordan et al. 2002; Bergmiller et al. 2012; Luo et al. 2015), define core cellular processes (Wang et al. 2015), identify potential drug targets in pathogens and tumors (Roemer et al. 2003; Behan et al. 2019), and are the starting point to determine minimal genomes (Juhas et al. 2011; Hutchison et al. 2016). The fraction of essential genes within a genome reflects its complexity and redundancy, and anticorrelates with the number of encoded genes (Rancati et al. 2018). For instance, 80% of 482 genes in *M. genitalium* (Glass et al. 2006), 18% of ∼6,000 genes in *S. cerevisiae* (Giaever et al. 2002), and only 10% of the ∼20,000 genes in human cell lines (Blomen et al. 2015; Hart et al. 2015; Wang et al. 2015) are essential for viability. Essential genes tend to code for protein complex members (Dezso et al. 2003; Hart et al. 2007), play central roles in genetic networks (Costanzo et al. 2010), have few duplicates (Giaever et al. 2002), and share other properties (Deng et al. 2011; Hart et al. 2015) that differentiate them from non-essential genes, enabling their prediction (Hwang et al. 2009; Lloyd et al. 2015; Zhang et al. 2016). Although gene essentiality is significantly conserved, essentiality changes are frequent across species and even between individuals. For instance, 17% of the 1:1 orthologs between *S. cerevisiae* and *S. pombe* have different essentiality (Kim et al. 2010). Also, 57 genes differ in essentiality between two closely related *S. cerevisiae* strains (Dowell et al. 2010), and a systematic analysis of 324 cancer cell lines from 30 cancer types found that only ∼40% of essential genes were shared across cell lines (Behan et al. 2019). Thus, essentiality is not a static property, and changes in the genetic background can change the essentiality of a gene (Rancati et al. 2018).

Recently, we and others have systematically identified essential genes that are non-essential (i.e. dispensable essential genes) in the presence of suppressor mutations (i.e. the genetic changes enabling the bypass of gene essentiality) in *S. cerevisiae* (Liu et al. 2015; van Leeuwen et al. 2020) and *S. pombe* (J. Li et al. 2019; Takeda et al. 2019). Both dispensable essential genes (DEGs) and their bypass suppressors exhibit specific features that differentiate them from other essential genes (i.e. core essential genes) and passenger mutations (i.e. randomly acquired mutations without an effect on fitness). For instance, DEGs are more likely to have paralogs, to be absent in other species, and encode members of smaller protein complexes compared to core essential genes (Liu et al. 2015; van Leeuwen et al. 2020), whereas suppressor genes tend to be functionally related to the DEG (van Leeuwen et al. 2020). We previously exploited the specific properties of these genes for their accurate prediction (van Leeuwen et al. 2020).

Identification of the suppressor genes responsible for bypassing the requirement for the essential gene is important to dissect the function of both genes (van Leeuwen et al. 2016), to expose the genetic architecture of phenotypic traits (Mackay 2014; Wei et al. 2014), and to understand drug resistance mechanisms (Woodford and Ellington 2007). Suppressor mutations could also explain the presence of presumably highly detrimental genetic variants in natural populations (Jordan et al. 2015; Narasimhan et al. 2015; Chen et al. 2016). For instance, highly penetrant disease-associated mutations are sometimes present in healthy individuals (Chen et al. 2016), and human pathogenic variants can be fixed in other mammalian species without obvious deleterious consequences (Jordan et al. 2015). However, whether suppression interactions identified in laboratory strains are relevant in natural evolutionary landscapes and could explain the presence of deleterious genetic variants in populations remains an open question.

Here, we compiled a comprehensive set of DEGs in *S. cerevisiae* identified across different studies to exhaustively compare their properties to core essential and non-essential genes, with a particular focus on phylogenetic features. We integrated bypass suppressor genes into an interaction network with DEGs to identify prevalent interaction motifs and to analyze the relationship of bypass suppression pairs in other species. This work presents a systematic characterization of DEGs and explores how their interactions with suppressors reflect evolution in natural populations.

## RESULTS

### Dispensable essential gene datasets

We compiled a comprehensive list of dispensable essential genes (DEGs) in *S. cerevisiae* from two large-scale studies (Liu et al. 2015; van Leeuwen et al. 2020) and from individual cases described in the literature (van Leeuwen et al. 2020)(**Figure 1A**). In total, 205 dispensable essential genes had been identified, representing ∼20% of all tested essential genes (**Figure 1B**). To determine whether the datasets could be merged, we compared various properties of the DEGs described in each dataset. The DEGs identified in the three datasets overlapped significantly (p < 0.001). In all datasets, DEGs showed similar functional enrichments (**Figure S1A**), and were depleted for fundamental cellular processes like RNA processing or translation and enriched for more peripheral functions related to signaling or transport. Furthermore, protein complexes tended to be either completely dispensable or indispensable across datasets (**Figure S1B**). For instance, the combined dataset contained 14 protein complexes with only dispensable essential subunits (**Figure S1C**), significantly more than expected by chance (**Figure S1B**). DEGs were more likely than core essential genes to be non-essential in the closely related *S. cerevisiae* strain Sigma1278b (**Figure S1D**), and to be absent in the *S. cerevisiae* core pangenome (**Figure S1E**). Since the properties of the combined and individual datasets were similar, we used the combined dataset in the following analyses.

**Figure 1.**
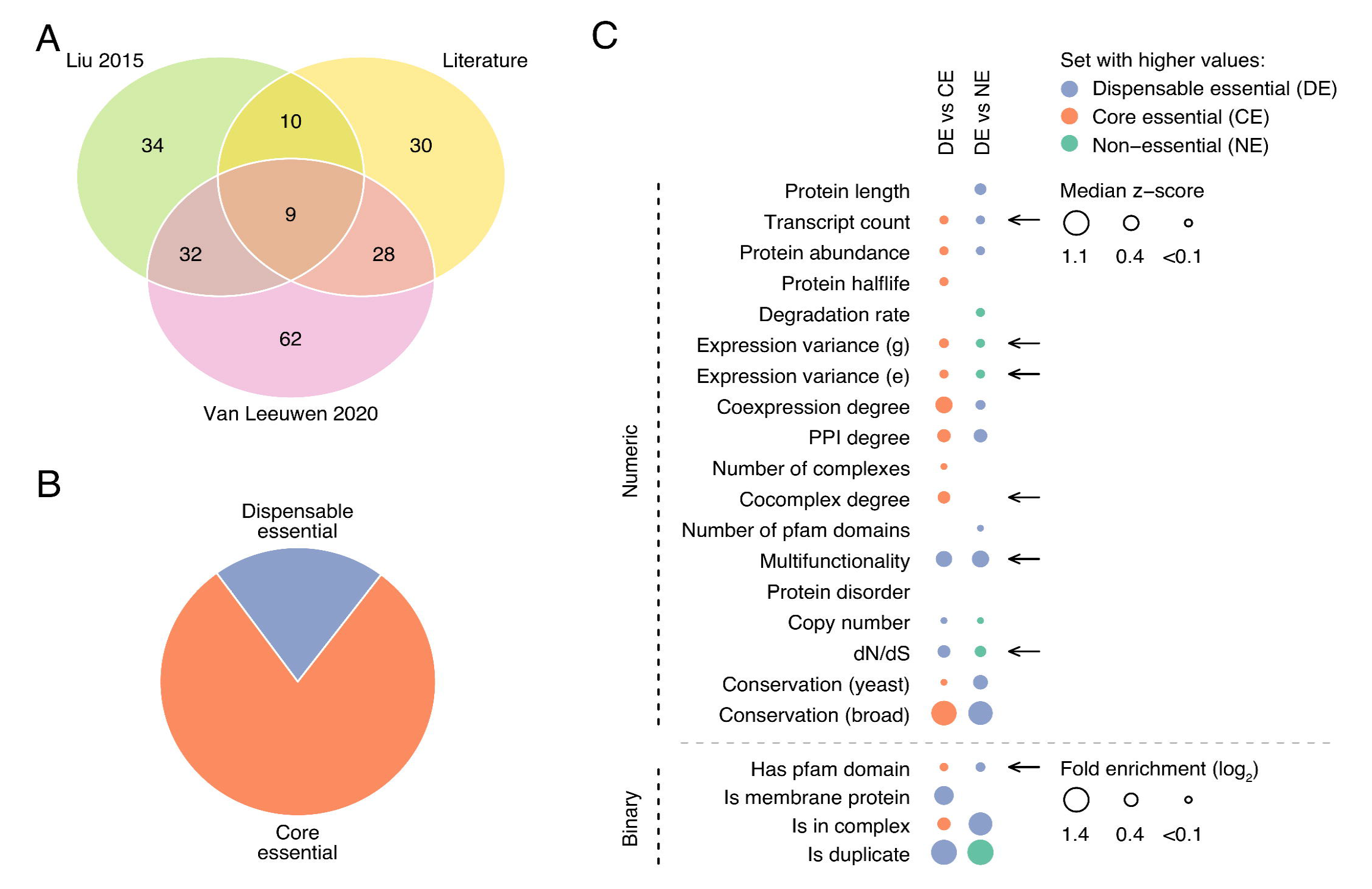
Properties of dispensable essential gene sets. **A**: Number of dispensable essential genes per individual dataset and their overlap. **B**: Fraction of dispensable and core essential genes in the combined dataset. **C**: Enrichment of dispensable essential vs core essential genes (left column) and vs non-essential genes (right column) for a panel of gene features. Top and bottom panels include numeric and binary features, respectively. Dot size is proportional to the fold enrichment and only enrichments with p-value < 0.05 are shown. The arrows indicate properties of dispensable essential genes not previously identified.

**Figure S1.**
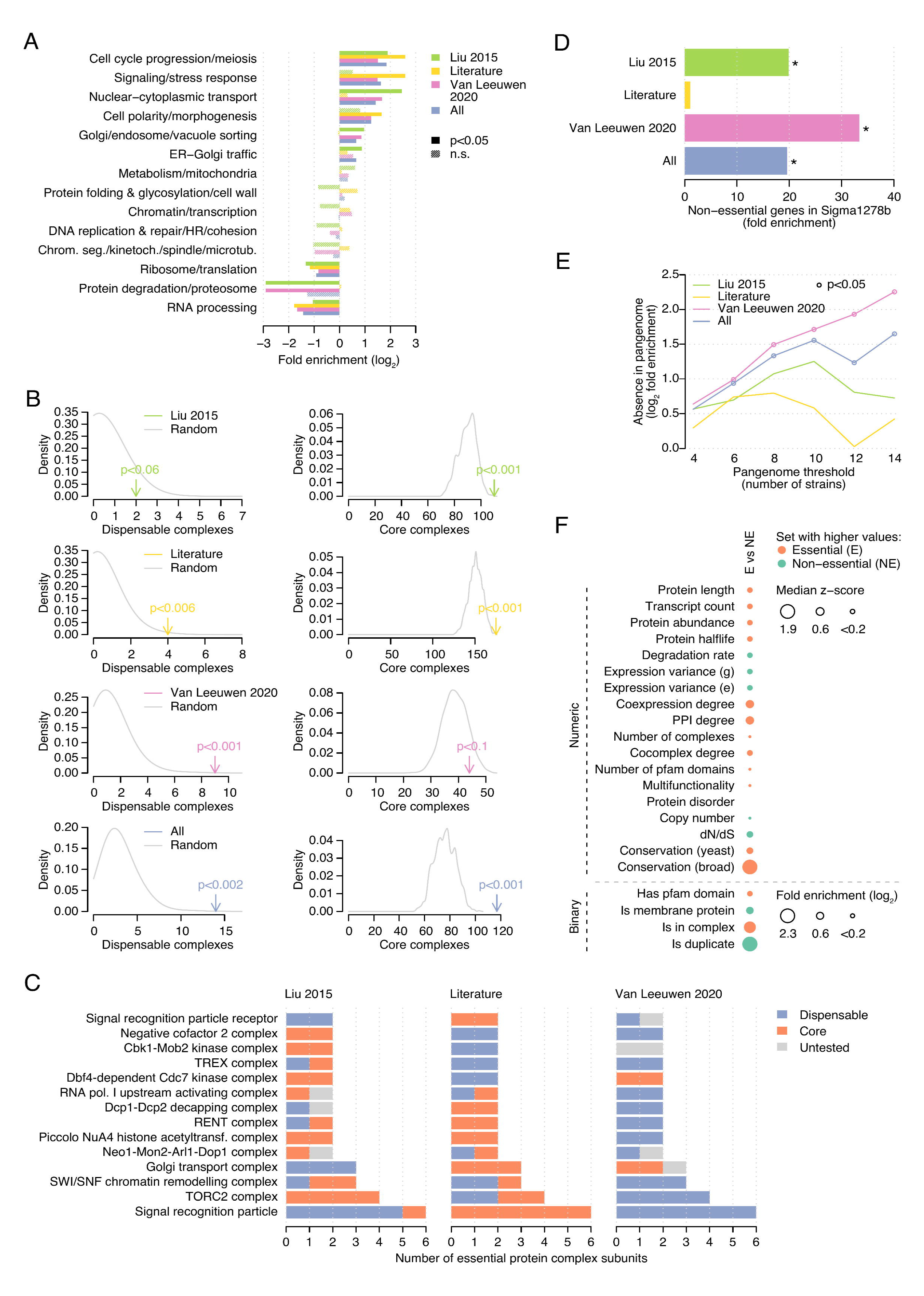
**A:** Functional enrichments of each dataset of dispensable essential genes. **B**: (left) Number of dispensable essential (i.e. all essential subunits are dispensable) and (right) core essential complexes (i.e. all essential subunits are core essential genes) for each dispensable gene set. The number of complexes in randomly selected gene sets of the same size are indicated in grey. **C**: The 14 complexes for which all essential genes are dispensable in the combined dataset. The subunits are colored by their dispensability in each individual dataset. Note that for the literature dataset we consider all essential genes as tested. **D**: Fold enrichment of each dataset of dispensable essential genes for non-essential genes in the Sigma1278b *S. cerevisiae* strain. **E**: Overlap between dispensable essential genes and missing genes in the core pangenome, defined at different thresholds. Fold enrichment with respect to the core essential genes is shown. **F**: Enrichment of non-essential genes vs essential genes for a panel of gene features. Left and right panels include binary and numeric features, respectively. Dot size is proportional to the fold enrichment and only enrichments with p-value < 0.05 are shown.

### Properties of dispensable essential genes

By querying an extensive panel of gene features, we compared the properties of dispensable and core essential genes. DEGs tended to exhibit more stable gene expression levels and lower transcript counts, to be less conserved across species, to have more gene duplicates and higher evolutionary rates, and to be coexpressed with fewer genes than core essential genes. The proteins encoded by DEGs tended to be more multifunctional, to lack structural domains, to localize to a membrane, to be absent from protein complexes, and to have fewer protein-protein interactions (PPIs), lower abundances, and shorter half-lives compared to those encoded by core essential genes (**Figure 1C**). Interestingly, the observed differences between dispensable and core essential genes resembled the differences between non-essential and essential genes (**Figure S1F**). Thus, we asked whether dispensable essential and non-essential genes shared the same properties, and found they comprised two different classes of genes with clearly distinct features (**Figure 1C**). Broadly, features of DEGs fell between those of core essential and non-essential genes, consistent with and extending previous findings in a smaller dataset (Liu et al. 2015).

### Phylogenetic analysis of dispensable essential genes

We further explored the differences in gene conservation between dispensable and core essential genes using the phylogeny of *S. cerevisiae*, starting with a large panel of sequenced *S. cerevisiae* strains (Peter et al. 2018). DEGs were more likely than core essential genes, but less than non-essential genes, to harbor deleterious mutations disrupting protein sequences (**Figure S2A**), to present higher nonsynonymous mutation rates (**Figure S2B**), and to show copy number loss (CNL) events in other *S. cerevisiae* strains (**Figure S2C**). To further investigate differences in the evolutionary pressure on dispensable essential and core essential genes, we analyzed essentiality data and orthology relationships in *Candida albicans*, *Schizosaccharomyces pombe*, and human cell lines (**Figure 2A, S2D, S2E**). Genes that were dispensable essential in *S. cerevisiae* were more often absent than core essential genes in each of the analyzed species (**Figure 2B**). We hypothesized that this bias could be caused by: i) genes specific to the *S. cerevisiae* phylogenetic branch and, thus, not present in their common ancestor, or ii) genes present in their common ancestor but lost in the phylogenetic branch of the analyzed species. To determine the contribution of each factor, we calculated the age of each *S. cerevisiae* gene by identifying the furthest species with an orthologous gene. DEGs were enriched for younger genes with respect to core essential genes (**Figure 2C**), particularly for genes with no ortholog in any other species (i.e. specific to *S. cerevisiae;* **Figure 2D**). Next, for each species we defined lost genes as those absent in that species but present in its common ancestor with *S. cerevisiae*. We found DEGs were more often lost in other species than core essential genes (**Figure 2E**). Thus, the absence of DEGs in other species can be explained both by genes specific to *S. cerevisiae* and by gene loss events in those species.

**Figure 2.**
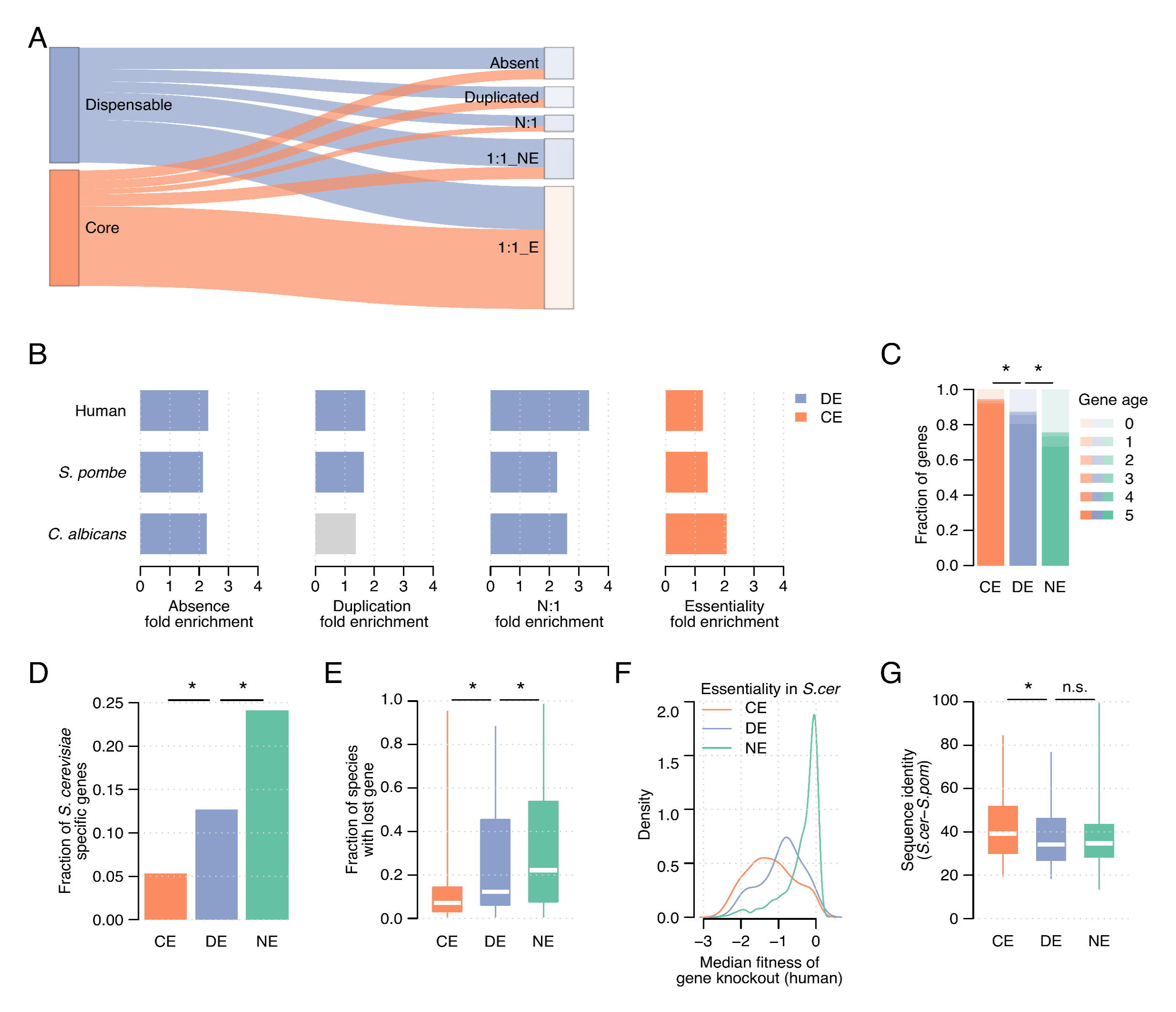
Phylogenetic analysis of dispensable essential genes. **A**: Orthology relationships in *S. pombe* of dispensable and core essential *S. cerevisiae* genes. The fraction of absent, duplicated, N:1, and essential and non-essential 1:1 orthologs is shown for each gene set. **B**: Fold enrichment of dispensable essential *S. cerevisiae* genes with respect to core essential genes for absence, duplication, N:1 relationships, and essential 1:1 orthologs in *S. pombe*, *C. albicans*, and human. Purple and orange bars identify significant enrichments for dispensable essential and core essential genes, respectively. Grey bars identify non-significant enrichments. **C**: Fraction of genes within each age group, ranging from zero (found only in *S. cerevisiae*) to five (found in the furthest ancestor), for the three sets of genes. **D**: Fraction of genes with age zero (*S. cerevisiae* specific) for each gene set. **E**: Fraction of gene loss events across species for each *S. cerevisiae* gene grouped by gene set. **F**: Median fitness per gene knockout across a panel of cancer cell lines. Genes are grouped by their essentiality in *S. cerevisiae*, and the density is shown. **G**: Protein sequence identity between gene products in *S. cerevisiae* and 1:1 orthologs in *S. pombe*. CE, core essential; DE, dispensable essential; NE, non-essential.

**Figure S2.**
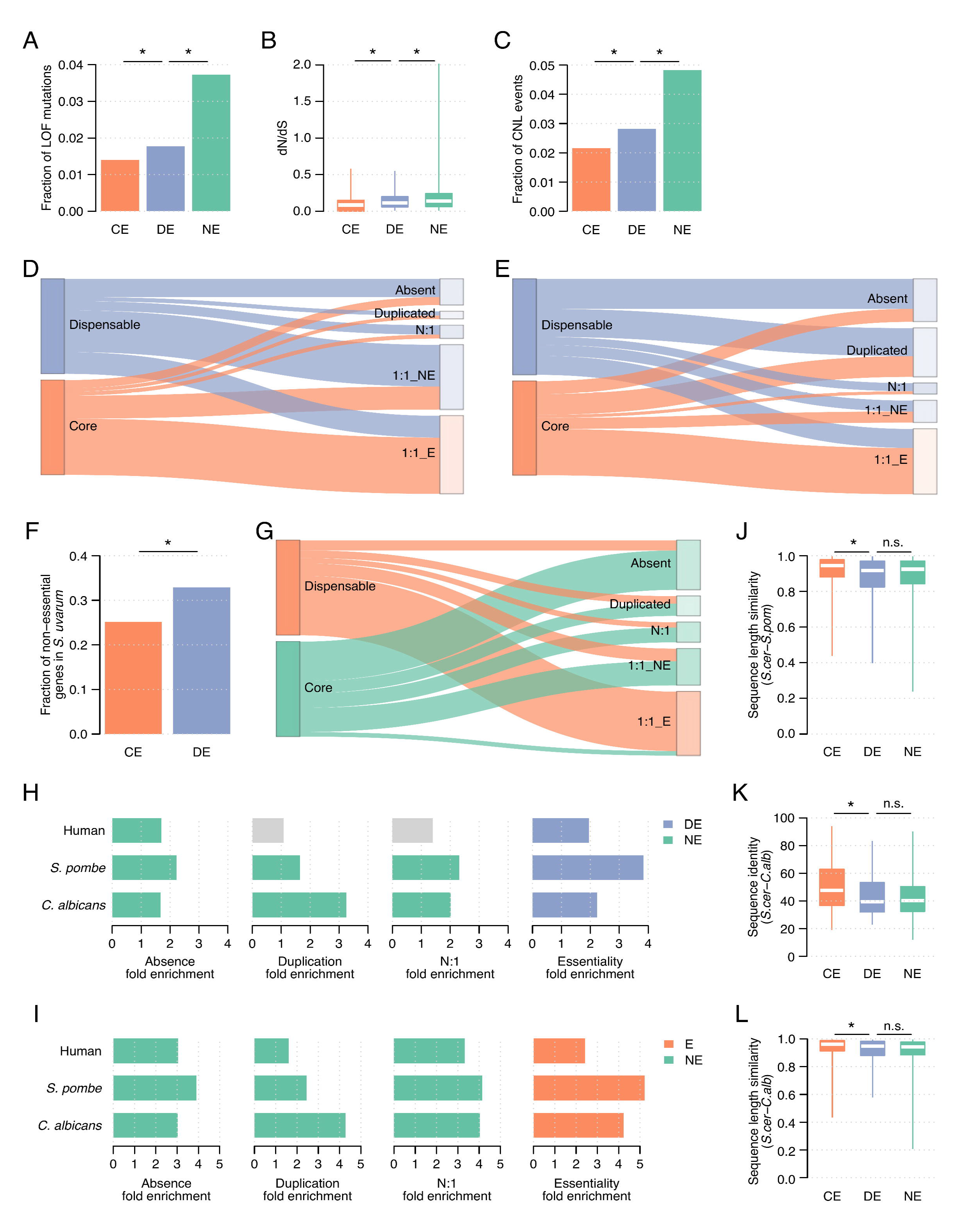
**A:** Fraction of loss-of-function mutations across *S. cerevisiae* strains for non-essential, dispensable essential, and core essential genes. **B**: Evolutionary rate (dN/dS) across *S. cerevisiae* strains for the three gene sets. **C**: Fraction of copy number loss events across *S. cerevisiae* strains for the three gene sets. **D**: Orthology relationships in *C. albicans* of dispensable essential and core essential genes. The fraction of absent, duplicated, N:1, and essential and non-essential 1:1 orthologs is shown for each gene set. **E**: Orthology relationships in human of dispensable essential and core essential genes. The fraction of absent, duplicated, N:1, and essential and non-essential 1:1 orthologs is shown for each gene set. **F**: Fraction of essential genes in *S. cerevisiae* that are non-essential in *S. uvarum*. **G**: Orthology relationships in *S. pombe* of non-essential and essential genes. The fraction of absent, duplicated, N:1, and essential and non-essential 1:1 orthologs is shown for each gene set. **H**: Fold enrichment of dispensable genes with respect to non-essential genes for absence, duplication, N:1 relationships, and essential 1:1 orthologs in *S. pombe*, *C. albicans*, and human. Blue and green bars identify significant enrichments for dispensable and non-essential genes, respectively. Grey bars identify non-significant enrichments. **I**: Fold enrichment of non-essential genes with respect to essential genes for absence, duplication, N:1 relationships, and essential 1:1 orthologs in *S. pombe*, *C. albicans*, and human. Green and orange bars identify significant enrichments for non-essential and essential genes, respectively. Grey bars identify non-significant enrichments. **J**: Ratio between the protein sequence length in *S. cerevisiae* and the 1:1 ortholog in *S. pombe*. The shorter length is divided by the longer one. **K**: Protein sequence identity between gene products in *S. cerevisiae* and 1:1 orthologs in *C. albicans*. **L**: Ratio between the protein sequence length in *S. cerevisiae* and the 1:1 ortholog in *C. albicans*. The shorter length is divided by the longer one. CE, core essential; DE, dispensable essential; NE, non-essential.

Furthermore, DEGs present in other species were more frequently duplicated and had more N:1 orthology relationships (**Figure 2B**) than core essential genes. For genes with a 1:1 ortholog in other species, DEG orthologs were more often non-essential than orthologs of core essential genes (**Figure 2B**), also in the closely related *S. uvarum* species (**Figure S2F**). Similarly, fitness data from a panel of 1,070 cancer cell lines (Meyers et al. 2017) revealed that knockout of DEG orthologs led to less severe proliferation defects than knockout of core essential gene orthologs (**Figure 2F**). Thus, genes that can be bypassed by genetic mutations in *S. cerevisiae* tend to be non-essential in other species. We show the comparison between essential and non-essential genes, and dispensable essential and non-essential genes to contextualize the observed differences (**Figure S2G-I**).

Finally, we compared sequences of *S. cerevisiae* proteins and their 1:1 orthologs in *S. pombe* and *C. albicans*. Gene products of DEGs had lower sequence identity and differed more in sequence length than core essential proteins (**Figure 2G, S2J-L**), in line with the dN/dS data (**Figure 1C, S2B**). Overall, orthology relationships, phenotypic changes, and sequence divergence reflect that the evolutionary pressure on DEGs is more lenient than on core essential genes but more strict than on non-essential genes.

### The bypass suppressor interaction network

Identification of the relevant genetic changes (i.e. suppressors) required to tolerate the deletion of an essential gene is key to interpreting the presence of deleterious genetic variants in natural populations. To improve our knowledge on the mechanisms of genetic suppression, we built an interaction network between DEGs and their bypass suppressors by combining data from our recent systematic study (van Leeuwen et al. 2020) and the literature (van Leeuwen et al. 2020). The two individual suppression interaction networks overlapped significantly (**Figure 3A**; p < 0.001) and were similarly enriched in functional associations (**Figure S3A**). The combined network included a total of 319 unique bypass suppression gene pairs, corresponding to 243 suppressors and 137 DEGs out of the 205 known DEGs. For the remaining DEGs (33% of the dataset), the suppressor variants have not been identified. Dispensable essential and suppressor genes tended to be functionally related (**Figure S3B**), particularly for close functional relationships like cocomplex or copathway membership (**Figure S3A**), and suppressors related to nuclear-cytoplasmic transport and transcription processes were more frequent than expected by chance (**Figure S3B**). For 50% and 26% of the dispensable genes, only LOF and GOF suppressors were isolated, respectively, and in 15% of the cases, both types of suppressors were identified (**Figure S3C**). For the remaining cases, the nature of the suppressor could not be determined. In agreement with the functional effect of the suppressor mutation, LOF alleles of genes in which LOF suppressor mutations had been identified showed more positive than negative genetic interactions with hypomorphic alleles of the corresponding DEG. Conversely, LOF alleles of suppressor genes in which a GOF mutation had been described had more negative than positive genetic interactions with hypomorphic mutants of the corresponding DEG (**Figure 3B**).

**Figure 3.**
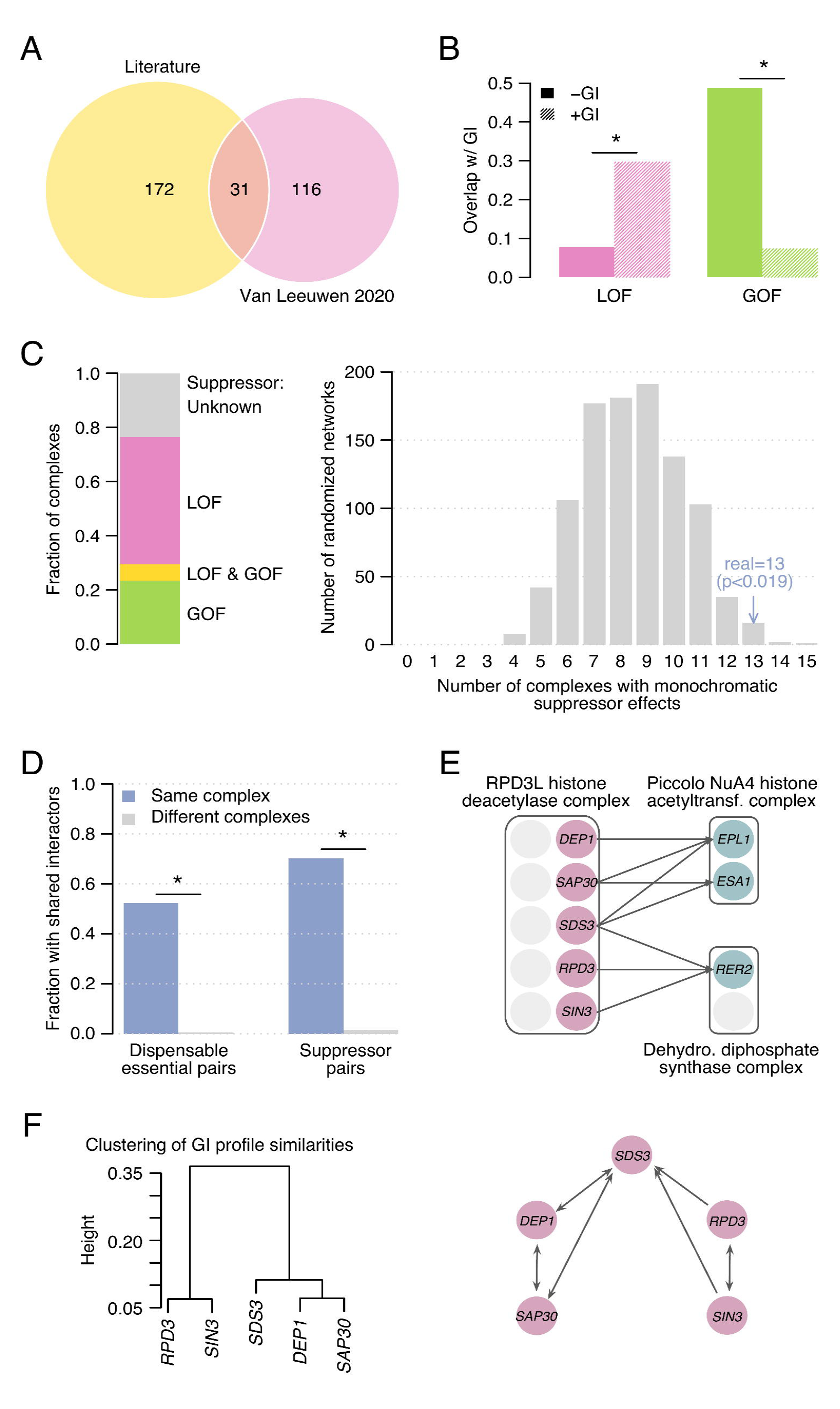
Bypass suppression interaction network. **A**: Number of bypass suppression gene pairs in each individual dataset and their overlap. **B**: Fraction of loss-of-function (LOF) and gain-of-function (GOF) bypass suppression pairs that overlap with negative and positive genetic interactions. **C**: (left) Fraction of monochromatic complexes in which all dispensable essential genes are suppressed by either loss-of-function (LOF) or gain-of-function (GOF) bypass suppressors. Only complexes with 2 or more dispensable essential subunits are shown. (right) Number of monochromatic complexes in the suppression bypass network (blue) and in 1,000 randomized networks (grey). **D**: Fraction of gene pairs encoding members of the same complex and of different complexes that share an interactor. Dispensable essential gene pairs are shown on the left, bypass suppressor gene pairs on the right. **E**: Interaction modularity of the bypass suppressor genes coding for members of the RPD3L histone deacetylase complex (CPX-1852). **F**: Genetic interaction profiles of the bypass suppressor genes in (E): (left) hierarchical clustering of the genetic interaction profiles; (right) network showing genetic interaction profile similarities above 0.2.

**Figure S3.**
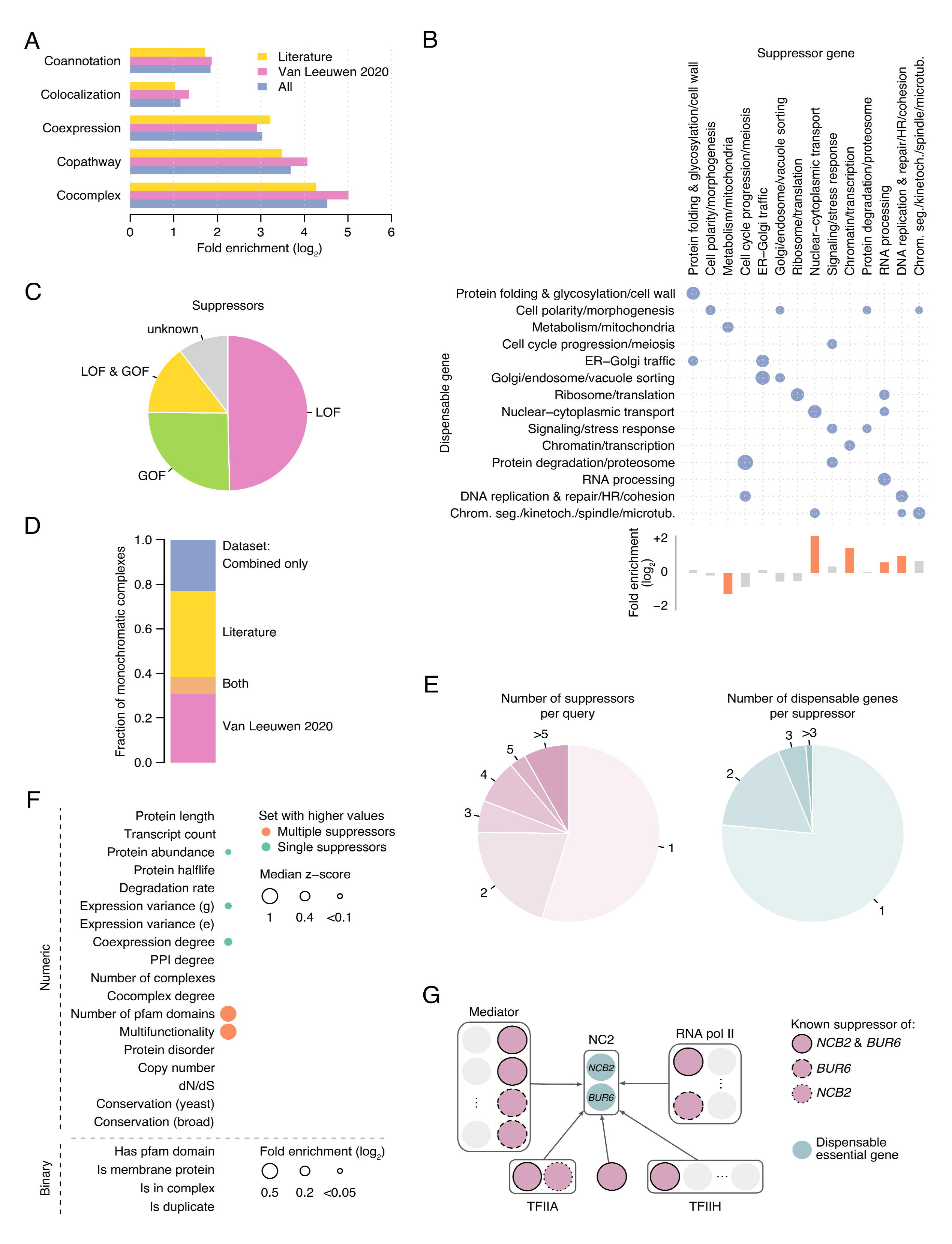
**A:** Functional enrichment for cocomplex, copathway, coexpression, colocalization, and GO co-annotation functional relationships of bypass suppression gene pairs in the combined and the two individual bypass suppression datasets. **B**: (central) Functional enrichment of interacting pairs in the bypass suppression network for 14 broad functional categories. Only significant enrichments are shown. Functional enrichments among dispensable essential (right) and suppressor genes (bottom) are also shown, in which orange and grey bars identify significant and non-significant associations, respectively. **C**: Fraction of dispensable essential genes with only loss-of-function suppressors (LOF), only gain-of-function suppressors (GOF), both loss-of-function and gain-of-function suppressors (LOF & GOF), and with suppressors of unknown type. **D**: Fraction of monochromatic complexes supported by each bypass suppression dataset. **E**: (left) Number of suppressors per dispensable essential gene. (right) Number of dispensable essential genes per suppressor. **F**: Enrichment of dispensable essential genes for which multiple suppressors have been described vs dispensable essential genes with a single identified suppressor for a panel of gene features. Dot size is proportional to the fold enrichment and only enrichments with p-value < 0.05 are shown. **G**: Interaction modularity of the bypass suppressor genes coding for members of the negative cofactor 2 complex (NC2, CPX-1662).

### Structure of the bypass suppression interaction network

Interaction density (i.e. the percentage of gene pairs with an interaction) of the bypass suppression network ranged from 0.007% to 0.96% depending on whether we considered all possible gene pairs or only pairs between the identified dispensable essential and suppressor genes, respectively. In spite of the sparsity of this network, several patterns emerge showing its structure and modularity. For instance, all DEGs in the same protein complex tended to interact with either GOF or LOF suppressors. These monochromatic interactions affected 13 out of 17 non-redundant protein complexes with at least two dispensable essential subunits in our dataset (**Figure 3C**; p < 0.05), suggesting similar suppression types apply for functionally related genes. Importantly, both individual suppression networks contributed to this result (**Figure S3D**), discarding the potential bias from specific hypothesis-driven experiments in the literature dataset. We analyzed the topology of the network and found that for 45% of the DEGs multiple suppressors had been described (**Figure S3E**). This set of genes exhibited specific features compared to DEGs for which only a single suppressor had been described (**Figure S3F**). For instance, DEGs with multiple identified suppressors tended to have higher multifunctionality and an increased number of structural domains, which suggest multiple different molecular mechanisms of suppression may exist for these DEGs. Suppressors were more specific than DEGs, and only 23% of them interacted with multiple genes (**Figure S3E**). Next, we explored the relationship between functional similarity and connectivity patterns. We found that genes in the same protein complex tended to have the same interactors: 52% of the DEGs encoding members of the same complex shared suppressor genes, and 70% of the suppressor genes encoding members of the same complex shared DEGs (**Figure 3D**), more than expected by chance (p < 0.05).

To illustrate the underlying modular structure of the bypass suppression interaction network, we explored the connectivity of *NCB2* and *BUR6*, both DEGs with known suppressors and the only two members of the negative cofactor 2 transcription regulator complex (ID CPX-1662 in the Complex Portal (Meldal et al. 2021)). *NCB2* and *BUR6* have seven and ten identified bypass suppressor genes, respectively, six of which are in common, again showing that functionally related DEGs tend to share suppressors (**Figure S3G**). Two of these common suppressors belong to the core Mediator complex that plays a role in the regulation of transcription (CPX-3226), showing that interactors of the same dispensable gene tend to be functionally related both to each other and to the DEG they are suppressing. The other four shared suppressor genes also affect transcription, and encode subunits of the transcription factor TFIIA complex (CPX-1633), the general transcription factor complex TFIIH (CPX-1659), and the DNA-directed RNA polymerase II complex (CPX-2662). Interestingly, the *NCB2* specific suppressor, *TOA2*, also belongs to TFIIA, and three of the four *BUR6* specific suppressors to RNA pol II or Mediator, further illustrating the modularity of the network. In another example (**Figure 3E**), members of the RPD3L histone deacetylase complex (CPX-1852) suppress two different protein complexes. *DEP1*, *SAP30*, and *SDS3* suppress the two essential subunits of piccolo NuA4 histone acetyltransferase complex (CPX-3185), whereas *RPD3*, *SIN3*, and *SDS3* interact with the Rer2 subunit of the dehydrodolichyl diphosphate synthase complex (CPX-162). This modularity in the suppression interaction pattern of RPD3L subunits is also observed in genome-wide genetic interaction patterns, which are more similar for RPD3L subunits that suppress the same query gene than for RPD3L subunits that suppress functionally diverse query genes (**Figure 3F**). These patterns suggest a functional modularity within the complex which is supported by its modeled structure (Sardiu et al. 2009).

### Mutational landscape of *S. cerevisiae* strains reflects bypass suppression relationships

We wondered if the genetic dependencies described in the suppression interaction network were reflected in the genomic variation present in natural populations. For that, we first evaluated if bypass suppression gene pairs in our network had simultaneous gene copy number changes across *S. cerevisiae* strains. For each gene pair and strain, we evaluated if the DEG showed a copy number loss (CNL) event (resembling the effect of a gene deletion) and the suppressor gene either a CNL or a copy number gain (CNG) event (equivalent to a LOF or a GOF effect, respectively). Interestingly, bypass suppression gene pairs with LOF and GOF suppressor mutations showed different preferences for co-occurring copy number changes. Bypass suppression gene pairs that involved a LOF suppressor mutation were enriched for co-loss of both dispensable essential and suppressor genes (**Figure 4A, S4A**). In contrast, cases with GOF suppressor mutations were enriched for events in which copy number loss of the dispensable essential was accompanied by a copy number gain of the suppressor gene (**Figure 4A, S4A**). Thus, when the DEG has a copy number loss in a natural strain, the functional effect of the bypass suppressor mutation (GOF or LOF) identifies the most likely copy number change of the suppressor gene in that same strain. Next, we asked whether deleterious coding mutations in DEGs and in identified bypass suppressor genes co-occurred in *S. cerevisiae* isolates. We only considered haploid strains so the deleterious effects of mutations would not be masked by other alleles. When considering only bypass suppression gene pairs in which the suppressor carried a LOF mutation, we found 18 cases in which both the DEG and the suppressor gene carried deleterious mutations in at least one of the haploid strains, significantly more than in randomized gene pairs (**Figure 4B**; p<0.05). As expected, we did not observe a similar enrichment in diploid strains (**Figure S4B**) nor for gene pairs involving GOF suppressor mutations (**Figure S4C**). Thus, the bypass suppression network mapped in a laboratory environment reflects evolutionary outcomes in natural *S. cerevisiae* strains.

**Figure 4.**
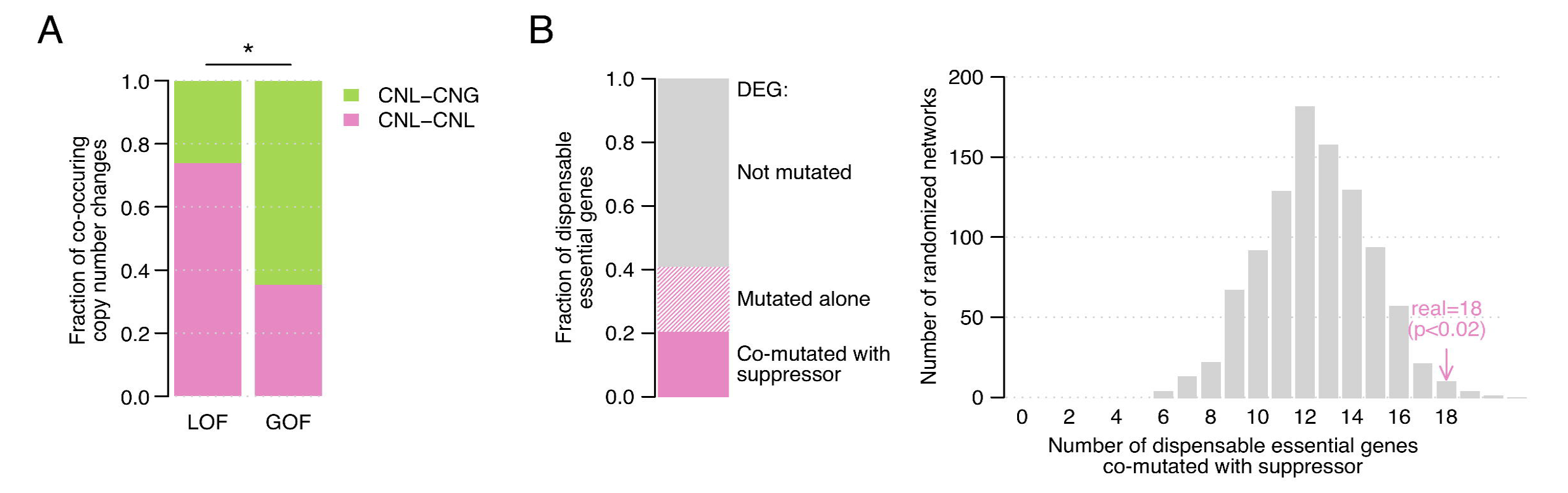
Co-occurring mutations in *S. cerevisiae* strains. **A**: Proportion of copy number co-loss and loss-gain (DEG-suppressor) events across a panel of *S. cerevisiae* strains for bypass suppression gene pairs in which the suppressor carried either a LOF or a GOF mutation. **B**: (left) Fraction of dispensable essential genes with no deleterious mutation across haploid *S. cerevisiae* strains, with a deleterious mutation in at least one of the strains but not co-occurring with deleterious mutations in any of its bypass suppressor genes, and with at least one strain in which it has a deleterious mutation co-occurring with a deleterious mutation in one of its known bypass suppressor genes. (right) Number of dispensable essential genes with a deleterious mutation in any of the haploid *S. cerevisiae* strains co-occurring with a deleterious variant in at least one of its known bypass suppressor genes using the bypass suppression network (pink) and a set of 1,000 randomized networks. In both analyses, only bypass suppression gene pairs with LOF suppressor mutations are considered.

**Figure S4.**
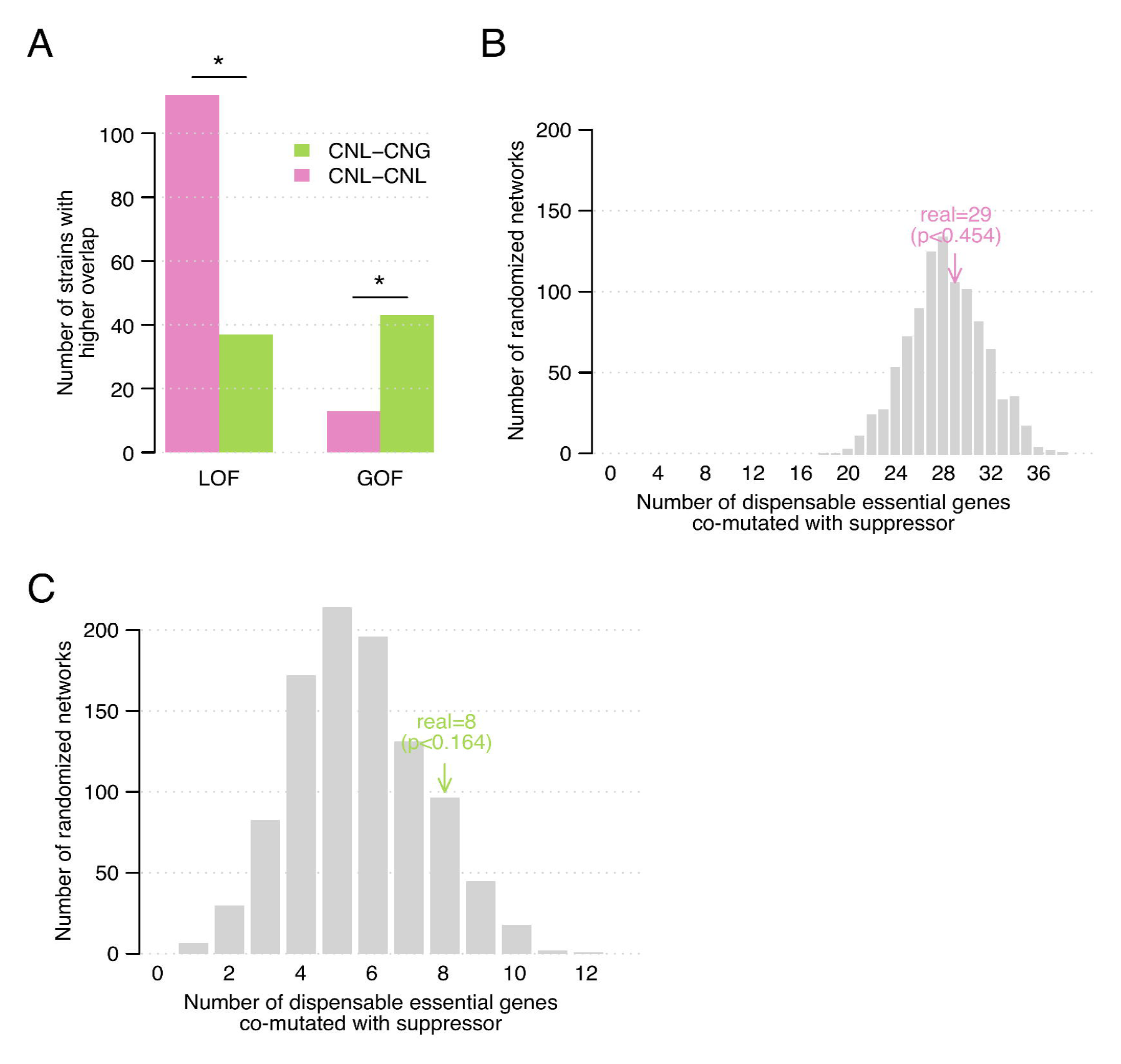
**A:** Number of *S. cerevisiae* strains in which bypass suppression gene pairs overlap more with copy number co-loss events than copy number loss-gain events (pink), and vice versa (green). Gene pairs are grouped by the bypass suppressor type: loss-of-function (LOF) and gain-of-function (GOF). **B**: Number of dispensable essential genes with a deleterious mutation in any of the diploid *S. cerevisiae* strains that co-occurs with a deleterious variant in at least one of its known bypass suppressor genes using the bypass suppression network (pink) and a set of 1,000 randomized networks. Only bypass suppression gene pairs with LOF suppressor mutations are considered. **C**: Number of dispensable essential genes with a deleterious mutation in any of the haploid *S. cerevisiae* strains that co-occurs with a deleterious variant in at least one of its known bypass suppressor genes using the bypass suppression network (green) and a set of 1,000 randomized networks. Only bypass suppression gene pairs with GOF suppressor mutations are considered.

### Co-occurrence of viability changes and fixed bypass suppressor mutations

We have shown that genes that are dispensable essential in *S. cerevisiae* are often non-essential in other species (**Figure 2B**). Differences in the genetic background in those species may be responsible for these changes in essentiality. Here, we hypothesized that the genetic changes that bypass the essentiality of a gene in *S. cerevisiae* should be reflected in the genome of species in which the gene is also dispensable (i.e. non-essential or absent). To test this, we evaluated whether changes in essentiality for DEGs in a given target species co-occurred with bypass suppressor mutations that were fixed in the target genome. Briefly, we considered as equivalent bypass mutations those that could reduce or increase the gene activity in the target species, for LOF and GOF suppressors, respectively (see Methods). Given that genome-scale essentiality data is scarce, we focused our analysis on *S. pombe*, for which high quality essentiality data is available for most genes (Harris et al. 2022).

We found that 67% (18/27) of the *S. cerevisiae* DEGs that are non-essential in *S. pombe* co-occurred with bypass suppressor mutations in that species, whereas this happened for only 26% (12/47) of the DEGs that were essential in *S. pombe* (2.6 fold enrichment; p < 0.05; **Figure 5A, 5B**). A similar trend (48%) was observed for *S. cerevisiae* DEGs that were absent (i.e. without an ortholog) in *S. pombe*, although this difference was not significant compared to the set of essential orthologs (**Figure 5B**). In order to increase the statistical power of our analyses, we combined the non-essential and absent genes in *S. pombe* into a single set and observed a clear difference with the essential orthologs (2.3 fold enrichment; p < 0.05).

**Figure 5.**
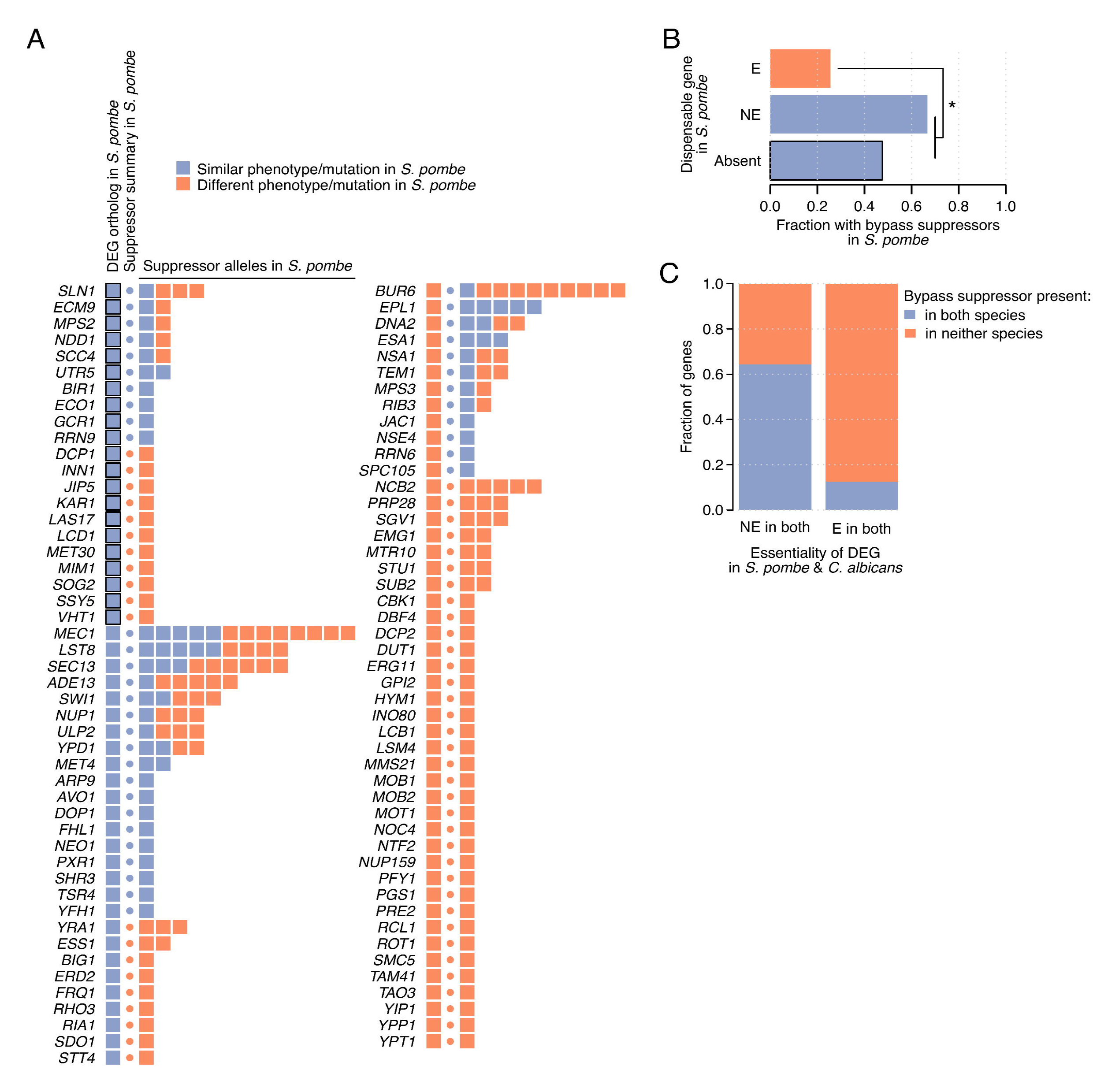
Changes in essentiality co-occur with bypass suppressor mutations. **A**: Dispensable essential *S. cerevisiae* genes without an ortholog or with a 1:1 ortholog in *S. pombe*, and their bypass suppressors. Color code reflects whether dispensable essential and bypass suppressor genes have similar phenotypes (i.e. absent or non-essential) and mutations, respectively, in *S. pombe* compared to the bypass suppression interactions identified in *S. cerevisiae*. The circle indicates, for each dispensable essential gene, whether any of the bypass suppressor mutations is present in *S. pombe*. **B**: Fraction of dispensable essential genes with at least one bypass suppressor mutation in the *S. pombe* genome. Dispensable essential genes are grouped by the phenotype of their 1:1 ortholog in *S. pombe* (E: essential; NE: non-essential; absent: without an ortholog). **C**: Fraction of dispensable essential genes with bypass suppressor mutations in both *S. pombe* and *C. albicans* or in neither of those species. Dispensable essential genes are grouped by the essentiality of their 1:1 orthologs in those species.

**Figure S5.**
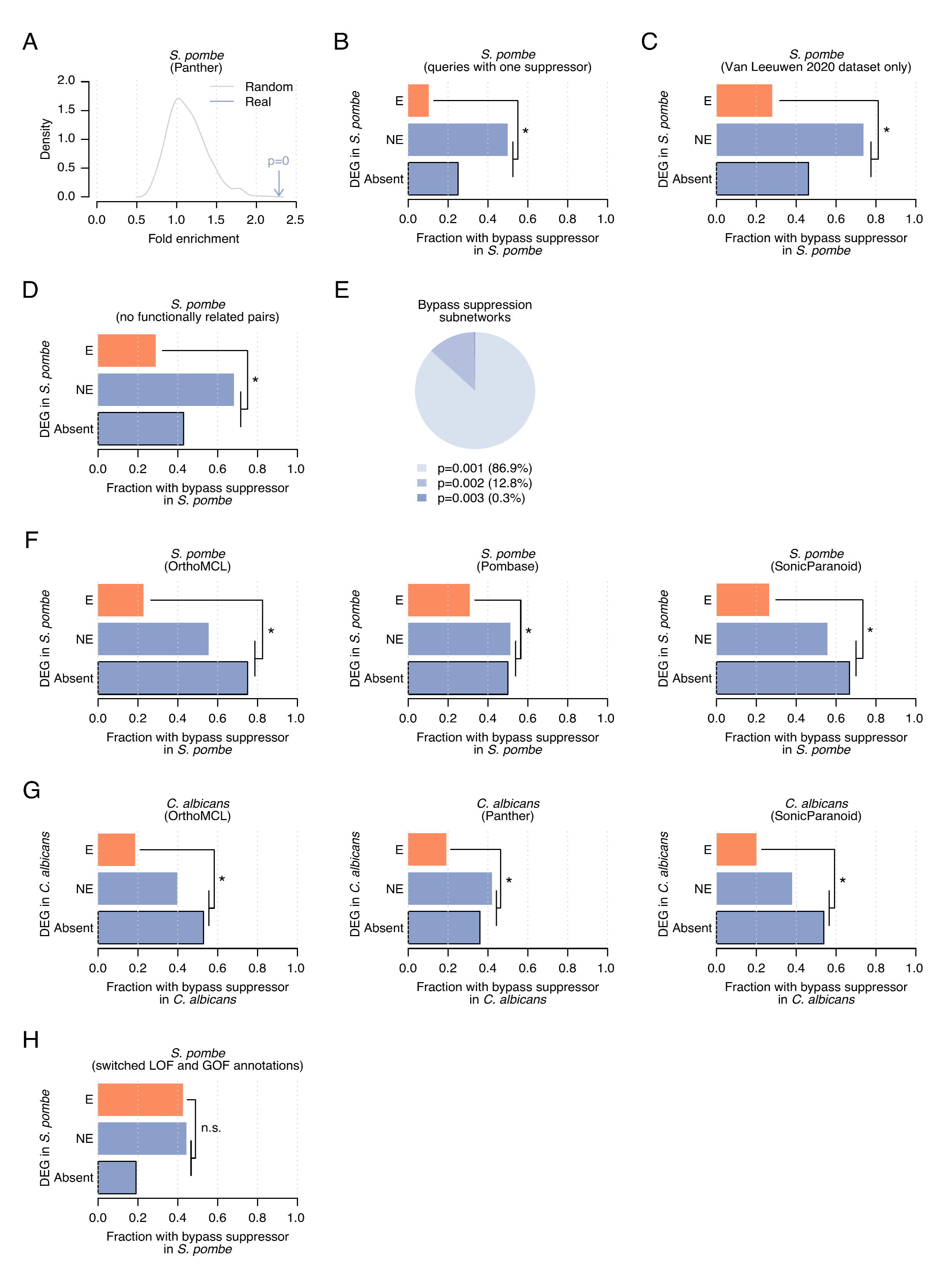
**A:** Presence of bypass suppressor mutations in the *S. pombe* genome for dispensable essential genes without an ortholog or with 1:1 orthologs that are non-essential in *S. pombe* vs dispensable essential genes with 1:1 orthologs that are essential. Fold enrichments for the suppression interaction network (blue) and 1,000 randomizations (grey) are shown. Blue squares with a black border identify dispensable essential genes without an ortholog in *S. pombe*. **B**: Like (5B) but considering only dispensable essential genes with a single bypass suppressor. **C**: Like (5B) but removing bypass suppression pairs from the literature. **D**: Like (5B) but removing bypass suppression pairs belonging to the same protein complex or pathway. **E**: Bypass suppression subnetworks grouped by the enrichment p-value of the co-occurrence of dispensable essential *S. cerevisiae* genes without orthologs or with 1:1 orthologs that were non-essential in *S. pombe*, and bypass suppressor mutations in the *S. pombe* genome. Each bypass suppression subnetwork has a different gene removed. **F**: Like (5B) but using different orthology mappings for *S. pombe*: OrthoMCL (left), Pombase (middle), and SonicParanoid (right). **G**: Like (5B) but using *C. albicans* orthology relationships and phenotype data: OrthoMCL (left), Panther (middle), and SonicParanoid (right). **H**: Like (5B) but with loss-of-function and gain-of-function annotations switched.

We controlled for potential biases to ensure the robustness of our observation (**Figure 5B**). We evaluated the effect of gene degree by generating 1,000 randomized bypass suppression networks while respecting the original topology (**Figure S5A**) and by considering only DEGs with a single known bypass suppressor (**Figure S5B**). Additionally, we removed bypass suppression interactions from the literature which may have been identified because of phylogenetic properties (**Figure S5C**), functionally related bypass suppression pairs which may be prone to present similar evolutionary patterns (**Figure S5D**), and every node in the network to discard dependence on a single gene (**Figure S5E**). Also, we applied three alternative orthology mappings (**Figure S5F**), and used essentiality annotations and orthology mappings from *C. albicans* (**Figure S5G**). In all these analyses, DEGs without orthologs or with non-essential orthologs more often co-occurred with bypass suppressor mutations than DEGs with essential orthologs. Conversely, switching LOF and GOF annotations resulted in a non-significant difference, as expected (**Figure S5H**).

Finally, we selected DEGs with 1:1 orthologs in both *S. pombe* and *C. albicans* and found that DEGs with non-essential orthologs in both species were more likely to have bypass suppressor mutations in those species than DEGs with essential orthologs (**Figure 5C**). In all, these analyses reveal that the relationship between DEGs and their bypass suppressor genes identified in *S. cerevisiae* is reflected in the gene essentiality and mutational space of other species.

## DISCUSSION

Differences between essential and non-essential genes have been widely characterized (**Figure S1F, S2G, S2I**) and a myriad of machine learning algorithms have exploited this information for the successful prediction of gene essentiality (Hwang et al. 2009; Lloyd et al. 2015; Zhang et al. 2016). Recently, we and others have identified a subset of *S. cerevisiae* essential genes that become dispensable in the presence of specific genetic variants (Liu et al. 2015; van Leeuwen et al. 2020). Here, we have combined these datasets of dispensable essential genes (DEGs), after showing they exhibit similar properties (**Figure 1**), for the comprehensive characterization of these genes. While recapitulating previously reported features in smaller datasets, we have also revealed new properties of dispensable essential genes (**Figure 1C, 2**). These features can be incorporated in existing methods for the prediction of essential gene dispensability (van Leeuwen et al. 2020). Since properties of DEGs are highly conserved (van Leeuwen et al. 2020), predictions could potentially target other species. Though the differences between dispensable essential and core essential genes resemble the differences between essential and non-essential genes (**Figure 1C, S1F, 2B, S2I**), dispensable essential and non-essential genes also make up two clearly distinct groups (**Figure 1C, S2H**). Thus, in contrast to the classical binary classification of genes based on their essentiality, three different sets of genes exist with specific properties that distinguish them from each other: non-essential, dispensable essential, and core essential genes, as was also previously suggested (Liu et al. 2015).

Importantly, we presented extensive evidence of the distinct evolutionary pressure exerted on these gene sets by performing phylogenetic analyses spanning very different evolutionary time-scales (**Figure 2, S2**), further expanding previous observations (Liu et al. 2015; van Leeuwen et al. 2020). The observed differences in conservation of dispensable essential compared to core essential *S. cerevisiae* genes in *S. uvarum*, *C. albicans*, *S. pombe*, and even human, which diverged from *S. cerevisiae* ∼1 billion years ago, reflect the substantial evolutionary footprint of essential gene dispensability.

For a better characterization of the mechanisms associated with the tolerance of highly deleterious mutations, we integrated data from multiple studies to build a bypass suppression interaction network between DEGs and their suppressors. Several properties emerged demonstrating the modularity and structure of the bypass suppression network. Complexes tended to be either composed of only dispensable essential subunits or of only core essential subunits (**Figure S1B**), mirroring the essentiality composition bias previously described (Hart et al. 2007) and the functional modularity that complexes encapsulate. Dispensable essentiality, thus, would be a modular feature of protein complexes (J. Li et al. 2019), as is essentiality. Also, protein complexes exhibited monochromaticity of suppressor type (**Figure 3C**), with members of the same complex being all suppressed by either LOF or GOF mutations. Last, members of the same complex exhibited interaction coherence, with cocomplexed DEGs sharing suppressors and cocomplexed suppressor genes interacting with the same DEGs (**Figure 3D**), as illustrated in **Figure 3E** and **S3G**. All these observations expose the inherent modularity of the bypass suppression network and suggest that similar suppression mechanisms apply for functionally related genes, which can lead to the identification of new dispensable essential and suppressor genes. Certainly, network modularity is not restricted to the bypass suppression network, and it is in fact a hallmark of a global genetic interaction network (Costanzo et al. 2016), but it is particularly relevant here given its directionality, small size, and low interaction density, reflecting the strong functional relationships bypass suppression interactions encapsulate.

The potential role of genetic suppression in explaining the existence of deleterious variants among natural populations (Chen et al. 2016) is still not fully understood. To address this knowledge gap, we evaluated how bypass suppression gene pairs reflected simultaneous genomic changes across evolution. Remarkably, we found co-occurrence of copy number changes and deleterious mutations in both the dispensable essential and the suppressor genes across *S. cerevisiae* strains. Furthermore, *S. cerevisiae* DEGs that were absent or non-essential in *S. pombe* were more likely to co-occur with a bypass suppressor mutation in the *S. pombe* genome than DEGs that were essential in *S. pombe* (**Figure 5**). These results suggest that within- and across-species genetic variability can follow the same evolutionary paths as spontaneous mutations in a laboratory environment, illustrating the constraints genetic networks may impose on evolutionary trajectories.

## CONCLUSION

We compiled a comprehensive list of dispensable essential genes across different studies which enabled the identification of new features of these genes. Phylogenetic analyses across *S. cerevisiae* strains and *S. pombe*, *C. albicans*, and human illustrated the more lenient evolutionary pressure affecting dispensable essential genes compared to core essential genes. The interaction network between dispensable essential and bypass suppressor genes exhibited a strong functional modularity. Importantly, the mutational landscape of *S. cerevisiae* strains reflected the bypass suppression relationships, with natural variants co-occurring in dispensable essential and bypass suppressor genes. Integration of phenotypic data from other species revealed that changes in essentiality across species also co-occur with the presence of fixed bypass suppressor mutations. Overall, our study provides an in-depth characterization of dispensable essential genes and unveils how bypass suppression relationships reflect on the evolutionary landscape.

## METHODS

### Dispensable essential gene analyses

#### Dispensable essential gene datasets

We retrieved DEGs in *S. cerevisiae* from two systematic experimental datasets (Liu et al. 2015; van Leeuwen et al. 2020) and from a study that compiled data from the literature (van Leeuwen et al. 2020). The set of tested genes are explicitly mentioned in the systematic studies, whereas for the literature set they are unknown and, therefore, we used all essential genes in *S. cerevisiae*. The combined dataset contained the DEGs identified in any of the three individual datasets. As tested genes, we considered all tested genes in the systematic studies and the dispensable genes identified in the literature set. We randomly generated 1,000 sets of genes of the same sizes as the individual datasets, sampling from the corresponding set of tested genes.

We calculated the overlap between the different datasets by counting the number of dispensable genes found across two and three datasets. We repeated the same process in the randomly generated datasets to derive empirical p-values.

#### Essentiality data

In our analyses, we used essentiality data from *S. cerevisiae* (van Leeuwen et al. 2020), *S. uvarum* (Sanchez et al. 2019), *C. albicans* (Segal et al. 2018), *S. pombe* (downloaded in November 2021 from Pombase (Harris et al. 2022)), and human cell lines (Hart et al. 2015). We considered human essential genes those that were required for viability in at least three of the five cell lines tested. In *C. albicans*, genes with essentiality confidence scores above 0.5 were classified as essential, and the remaining genes as non-essential.

#### Orthology mappings

We used PantherDB 16.1 (Mi et al. 2021) to identify orthology relationships. When indicated, we also used OrthoMCL (Li et al. 2003), SonicParanoid (Cosentino and Iwasaki 2019), based on the popular Inparanoid (Sonnhammer and Östlund 2015), and Pombase (Wood et al. 2012) orthology mappings.

#### Functional enrichment of dispensable essential genes

For each DEG set and functional class, we calculated the fold enrichment as the fraction of DEGs annotated to that functional class with respect to the corresponding fraction of core essential genes. Statistical significance was calculated with two-sided Fisher’s exact tests.

#### Enrichment for non-essential genes in the Sigma1278b strain

For each dispensable gene set, we calculated the fold enrichment as the ratio of DEGs identified as non-essential in the Sigma1278b strain divided by the analogous ratio of core essential genes. P-values were calculated using two-sided Fisher’s exact tests.

#### Complex dispensability bias

For each DEG set, we counted the number of complexes (Meldal et al. 2021) in which all essential subunits were identified either as dispensable or core essential genes. We repeated the same process using the randomly generated datasets to derive empirical p-values.

#### Properties of dispensable essential genes

We queried a panel of gene features previously defined (van Leeuwen et al. 2020). For numerical features, values of DEGs were z-score normalized using the median and standard deviation of the core essential genes. Dot size in plots is proportional to the median z-score value. We calculated the statistical significance by means of Mann-Whitney U tests. Only dots of significant differences (two-sided p-value < 0.05) are shown. For boolean features, we calculated the fold enrichment as the ratio of DEGs with that feature divided by the equivalent ratio of core essential genes. We calculated the p-values with Fisher’s exact tests. We followed the same approach to characterize: i) dispensable essential vs non-essential genes; ii) essential vs non-essential genes; iii) DEGs with multiple suppressors vs single suppressors.

#### Analyses on *S. cerevisiae* strains

We downloaded gene presence/absence data for a large panel of *S. cerevisiae* strains (G. Li et al. 2019) and defined several core pangenome gene sets at different stringency levels (see x-axis in **Figure S1E**). For instance, a threshold of ten identifies the core pangome composed of all genes absent only in ten strains or less. For each DEG dataset, we calculated the fraction of DEGs missing from the pangenome and the corresponding fraction for core essential genes, from which we calculated the fold enrichment. P-values were calculated with two-sided Fisher’s exact tests.

We retrieved precomputed loss-of-function data for *S. cerevisiae* strains (Peter et al. 2018) from http://1002genomes.u-strasbg.fr/files/, including frameshift mutations and missense mutations predicted to be deleterious by SIFT (Ng and Henikoff 2001). We calculated the number of strains in which these mutations affected each gene and aggregated the results per gene set (i.e, dispensable essentials, core essentials, and non-essentials). P-values were calculated using Fisher’s exact tests.

For each strain, we counted the genes affected by copy number loss (CNL) events in a panel of *S. cerevisiae* strains (Peter et al. 2018) and aggregated the result per gene set. P-values were calculated using Fisher’s exact tests. Then, we grouped the strains by their number of CNL events, which we used as a measure of distance to the reference strain, in three sets of equal size. For each set of strains, we calculated the proportion of CNL events that corresponded to each of the gene sets. P-values were calculated by Fisher’s exact tests comparing the fraction of CNL events corresponding to DEGs across strain sets. Finally, we retrieved dN/dS data for the same panel of *S. cerevisiae* strains and grouped them by gene set. P-values were calculated using Mann-Whitney U tests.

#### Orthology relationships of dispensable essential genes

For each gene, we calculated its orthology relationships in *C. albicans*, *S. pombe*, and human. Specifically, we considered gene absence, gene duplication (including 1:N and N:M orthology relationship), N:1 relationships, and 1:1 orthologs. For 1:1 orthologs, we evaluated the essentiality in the target species. For each species and property, the fold enrichment was calculated as the fraction of DEGs with respect to the fraction of core essential genes with that property. P-values were calculated by Fisher’s exact tests. We used the same approach to compare dispensable essential to non-essential genes, and non-essential to essential genes.

#### Gene age

For each gene, we calculated its age by identifying the farthest species from *S. cerevisiae* with a present ortholog. We used orthology relationships for 98 species from PantherDB (Mi et al. 2021). The phylogenetic tree to calculate species relationships was downloaded from Uniprot (UniProt Consortium 2021), and for each species we calculated the distance to *S. cerevisiae* as the number of main branches separating them. Thus, genes with age 0 are specific to *S. cerevisiae* and not present in any other of the 98 species, whereas age 5 corresponds to genes present in the most distantly related species. We grouped gene ages for each gene set (core, dispensable, and non-essentials) and calculated p-values with Mann-Whitney U tests.

#### Gene loss

For each gene of age X, we calculated the fraction of species closer to *S. cerevisiae* (distance < X) in the phylogenetic tree with that gene absent from their genome. For instance, for a given gene of age 3, we calculated the fraction of species at distance 1 or 2 to *S. cerevisiae* with the gene of interest absent. We aggregated data for each gene set (core essentials, dispensable essentials, and non-essentials) and calculated p-values by means of Mann-Whitney U tests. Also, we specifically evaluated gene loss taking place only in *S. pombe* and *C. albicans*, by considering genes with ages higher than their distance to *S. cerevisiae* (i.e. genes found in any of their common ancestors). P-values were calculated using Fisher’s exact tests.

#### Cancer cell lines

We used fitness data from genome-scale CRISPR-Cas9 knockout screens in 1,070 cancer cell lines from DepMap (Meyers et al. 2017). For each gene, we calculated the median effect of gene knockout on cell proliferation and the associated standard deviation across all cell lines. P-values were calculated using Mann-Whitney U tests.

#### Sequence analysis

For all 1:1 ortholog pairs between *S. cerevisiae* and *S. pombe*, we calculated their protein sequence identity. Sequence length similarity was calculated as the length ratio between the shortest of the sequences with respect to the longest. Thus, values closer to 1 describe sequence pairs of similar length, whereas values closer to 0 correspond to sequences of very different lengths. P-values were calculated using Mann-Whitney U tests. We followed the same approach to compare *S. cerevisiae* and *C. albicans* sequences.

### Suppression network analyses

#### Interaction data

We combined suppression interactions from our recent study (van Leeuwen et al. 2020) with interactions found in the literature (van Leeuwen et al. 2016) including only deletions of essential genes suppressed in standard conditions. We generated 1,000 randomized networks respecting the topology (i.e. maintaining the total number of connections of each gene) using the BiRewire R package (Iorio et al. 2016). We calculated the number of bypass suppression pairs present in both datasets and compared that value to the number of overlapping pairs in randomized networks to derive an empirical p-value.

#### Functional overlaps

We calculated the fraction of bypass suppression gene pairs that coded for proteins localized to the same subcellular compartment (Huh et al. 2003), had MEFIT (Huttenhower et al. 2006) coexpression scores above 1.0, were annotated to the same biological process GO term (Myers et al. 2006; Costanzo et al. 2016), and coded for members of the same complex (Meldal et al. 2021) and molecular pathway (Kanehisa et al. 2016). We repeated this calculation with the non-interacting gene pairs in the bypass suppression network, and derived fold enrichments and p-values using Fisher’s exact tests. We applied this approach to the individual and the combined datasets.

#### Complex monochromaticity by suppression mode

We selected a non-redundant set of 17 protein complexes with at least two dispensable essential subunits in the bypass suppression network. We only kept one representative complex when several complexes had the same set of DEGs. For each complex, we calculated if all dispensable essential subunits could be suppressed by the same suppressor mode (LOF or GOF). Note that in one complex, all subunits could be suppressed by LOF suppressors but also by GOF suppressors (indicated by “LOF & GOF” in the panel). We counted all complexes with this monochromaticity in suppression mode and compared that value to the number of monochromatic complexes in a set of 1,000 randomized bypass suppression networks to derive an empirical p-value. We applied the same approach to the two individual suppression networks to discard a bias in the literature dataset.

#### Network modularity based on cocomplex relationships

We counted the number of DEGs within the same protein complex (Meldal et al. 2021) that shared at least one suppressor. We repeated the same calculation using pairs of DEGs belonging to different complexes to derive a fold enrichment and a p-value calculated with a Fisher’s exact test. We followed the same approach querying for interactors of bypass suppressors instead of the interactors of DEGs.

#### Functional preferences

We annotated the dispensable essential and suppressor genes in the network using 14 broad functional classes (Costanzo et al. 2016). We then calculated the number of bypass suppression gene pairs within each pair of classes and repeated the process in randomized bypass suppression networks to derive empirical p-values. We used the median values of the randomized set to calculate the fold enrichments. Only fold enrichments of significant associations are shown.

We calculated fold enrichments for suppressors as the fraction of those genes in each class with respect to the corresponding fraction of background genes, from which we calculated a p-value using Fisher’s exact test.

#### Agreement in copy number changes and suppression mode across *S. cerevisiae* strains

We defined copy number loss (CNL) and copy number gain (CNG) events as having a copy number below 1 or above 1, respectively, per haploid genome as defined by the 1011 genomes project (Peter et al. 2018). For each bypass suppression gene pair, we calculated the number of strains in which both genes had a CNL (i.e. co-loss events) and a CNL event for the DEG and CNG for the suppressor gene (i.e. loss-gain events). We disregarded 18 hypermutated strains with copy-number changes in >33% of the genes in the bypass suppression network, and aggregated co-loss and loss-gain events for all bypass suppression gene pairs after splitting pairs by their suppression mode (LOF or GOF). We repeated the same calculation with a background set of gene pairs composed of all possible DEG-suppressor pairs, after removing the gene pairs in the bypass suppression network. We compared the proportion of co-loss vs loss-gain events for the LOF and GOF bypass suppression pairs, for LOF bypass suppression pairs and background pairs, and for GOF bypass suppression pairs and background pairs. We calculated the statistical significance by two-sided Fisher’s exact tests. Finally, we counted the number of strains in which LOF bypass suppression pairs overlapped more often with co-loss events than with loss-gain events, and vice versa, after normalizing by the event counts of the background pairs. We repeated the same process with GOF bypass suppression pairs. We calculated the statistical significance with two-sided binomial tests.

#### Co-mutation in *S. cerevisaie* strains

We defined as deleterious mutations the missense mutations predicted as damaging by SIFT, indel mutations, and frameshift mutations. For each DEG, we retrieved the strains in which it had a deleterious mutation and checked if any of its bypass suppressor genes was also mutated in any of those strains. We counted the number of DEGs co-mutated in any strains with any of their suppressor genes, the number of dispensable genes mutated alone, and the number of dispensable genes not mutated in any strain. We repeated the same process using 1,000 randomized bypass suppression networks. We performed this calculation using: 1) LOF bypass suppression pairs and haploid strains; 2) LOF bypass suppression pairs and diploid strains; and 3) GOF bypass suppression pairs and haploid strains.

#### Phenotypic changes across species and presence of bypass suppressor mutations

We hypothesized that the relationships between DEGs and bypass suppressor mutations identified in *S. cerevisiae* should be reflected in the evolutionary landscape of other species. To test this hypothesis, we identified DEGs that were non-essential or absent in a given target species and evaluated if the bypass suppressor mutations were fixed in the given target genome. To determine if a bypass suppressor mutation was fixed in another species, we took into account the effect of the suppressor mutation on gene function. Briefly, for LOF suppressors, we evaluated if mutations in the target species would reduce the gene activity with respect to *S. cerevisiae*. Conversely, for GOF suppressors, we evaluated if the mutations would increase the gene activity.

First, we annotated the orthology relationship of each DEG in *S. pombe*. We only considered DEGs absent in *S. pombe* or with a 1:1 ortholog. For genes with 1:1 orthologs, we annotated the essentiality of the ortholog in that species. We also annotated the orthology relationships of bypass suppressor genes in *S. pombe*. For suppressors with 1:1 orthologs, we performed a sequence alignment between the protein sequences of both species.

We next describe the set of rules that we evaluated to identify cases with equivalent bypass mutations in *S. pombe*. Briefly, in LOF suppressors we looked for orthologs with decreased activity with respect to the *S. cerevisiae* gene, whereas in GOF suppressors, for orthologs with increased activity. The first set of rules was based on orthology relationships. We considered *S. pombe* to have a LOF bypass mutation if the suppressor gene was absent. Also, if it had an N:1 ortholog, which could be similar to a copy number decrease and, thus, a decrease in activity. Suppressors with more than one ortholog in *S. pombe* or with a 1:1 ortholog were considered non-equivalent LOF bypass mutations, since their copy number did not decrease. Conversely, we considered as GOF bypass mutations cases in which the suppressor gene had more than one ortholog, similar to increasing their copy number and their activity, and non-equivalent GOF bypass mutations cases in which there was a N:1, 1:1, or absent ortholog in *S. pombe*.

The second set of rules we used to evaluate equivalent mutations was based on protein sequences. We only considered frameshift, nonsense, and missense mutations of suppressor genes with 1:1 orthologs. For the rest of cases, only the orthology rules (see above) were applied. The position of the nonsense and frameshift suppressor mutations identifies the part of the protein that should remain functional. Functionality encoded beyond that residue is compromised. Thus, we considered 1:1 orthologs in *S. pombe* with a shorter sequence than the position of the nonsense or frameshift suppressor mutation as LOF bypass mutations. Conversely, we considered cases in which the ortholog sequence was equal or longer than the position of the nonsense or frameshift suppressor mutation as non-equivalent LOF bypass mutations. In cases with missense mutations, we performed a sequence alignment between the *S. cerevisiae* suppressor gene and its 1:1 ortholog in *S. pombe*. We considered the ortholog to have an equivalent LOF bypass mutation if the same mutated residue or a gap was found in the aligned mutated position of the ortholog sequence. If the aligned mutated residue was the same as in the wild-type *S. cerevisiae* sequence (i.e. unmutated), we considered the ortholog to have a non-equivalent LOF bypass mutation. Cases in which the aligned position had different residues in *S. pombe* (not the wild-type and not the suppressor mutation) could not be classified as either equivalent or non-equivalent LOF bypass mutations. For GOF suppressors with a missense mutation and a 1:1 ortholog in *S. pombe*, we also performed a sequence alignment between the suppressor gene and its 1:1 ortholog. We considered the ortholog to have an equivalent GOF bypass mutation if the same mutated residue was found in the aligned mutated position of the ortholog sequence. If the aligned mutated residue was the same as in the wild-type *S. cerevisiae* sequence (i.e. unmutated), we considered the ortholog to have a non-equivalent GOF bypass mutation. The rest of cases could not be classified as either equivalent or non-equivalent GOF bypass mutations. We also evaluated missense mutations of suppressors with unknown suppression mode that had 1:1 orthologs in *S. pombe*. Cases with the exact same mutation in the ortholog were classified as equivalent bypass mutations, whereas cases in which the residue did not change in the 1:1 ortholog were classified as non-equivalent bypass mutations. The remaining suppressors with unknown suppression mode were not evaluated. Importantly, in suppressor genes with a frameshift, nonsense, or missense mutation, and with a 1:1 ortholog in *S. pombe*, the sequence based assessment took precedence over the orthology based evaluation.

Finally, we considered a DEG to have an equivalent bypass suppressor in *S. pombe* if any of its suppressors satisfied that criteria. We grouped DEGs by their essentiality in *S. pombe*, expecting DEGs with equivalent phenotypes in *S. pombe* (i.e absent or 1:1 non-essential orthologs) to have equivalent bypass suppressors more often than DEGs with a 1:1 essential ortholog. We calculated the fraction of genes with equivalent bypass suppressors for both gene sets to derive a fold enrichment and the p-value with a one-sided Fisher’s exact test. We compared the fold enrichment of the bypass suppression network to a set of randomized bypass suppression networks, which we used to derive an empirical p-value.

We repeated the exact same process: i) using *C. albicans* sequences, orthology relationships, and essentiality annotations; ii) using orthoMCL (Li et al. 2003), SonicParanoid (Cosentino and Iwasaki 2019), and Pombase (Wood et al. 2012) as alternative orthology mappings; iii) considering only DEGs with a single bypass suppressor to control for the bias introduced by gene degree; iv) removing bypass suppression pairs from the literature which may have been potentially identified by phylogenetic approaches; v) removing cocomplex and copathway bypass suppression pairs which may be more prone to present similar phylogenetic patterns; vi) switching LOF and GOF annotations to demonstrate the specificity of our sets of rules; vii) removing every node in the network to discard dependence on a single gene.

## REFERENCES

Behan FM, Iorio F, Picco G, Gonçalves E, Beaver CM, Migliardi G, Santos R, Rao Y, Sassi F, Pinnelli M, et al. 2019. Prioritization of cancer therapeutic targets using CRISPR-Cas9 screens. Nature 568:511–516.

Bergmiller T, Ackermann M, Silander OK. 2012. Patterns of evolutionary conservation of essential genes correlate with their compensability. PLoS Genet. 8:e1002803.

Blomen VA, Májek P, Jae LT, Bigenzahn JW, Nieuwenhuis J, Staring J, Sacco R, van Diemen FR, Olk N, Stukalov A, et al. 2015. Gene essentiality and synthetic lethality in haploid human cells. Science 350:1092–1096.

Chen R, Shi L, Hakenberg J, Naughton B, Sklar P, Zhang J, Zhou H, Tian L, Prakash O, Lemire M, et al. 2016. Analysis of 589,306 genomes identifies individuals resilient to severe Mendelian childhood diseases. Nat. Biotechnol. 34:531–538.

Cosentino S, Iwasaki W. 2019. SonicParanoid: fast, accurate and easy orthology inference. Bioinformatics 35:149–151.

Costanzo M, Baryshnikova A, Bellay J, Kim Y, Spear ED, Sevier CS, Ding H, Koh JLY, Toufighi K, Mostafavi S, et al. 2010. The genetic landscape of a cell. Science 327:425–431.

Costanzo M, VanderSluis B, Koch EN, Baryshnikova A, Pons C, Tan G, Wang W, Usaj M, Hanchard J, Lee SD, et al. 2016. A global genetic interaction network maps a wiring diagram of cellular function. Science [Internet] 353. Available from: http://dx.doi.org/10.1126/science.aaf1420

Deng J, Deng L, Su S, Zhang M, Lin X, Wei L, Minai AA, Hassett DJ, Lu LJ. 2011. Investigating the predictability of essential genes across distantly related organisms using an integrative approach. Nucleic Acids Res. 39:795–807.

Dezso Z, Oltvai ZN, Barabási A-L. 2003. Bioinformatics analysis of experimentally determined protein complexes in the yeast Saccharomyces cerevisiae. Genome Res. 13:2450–2454.

Dowell RD, Ryan O, Jansen A, Cheung D, Agarwala S, Danford T, Bernstein DA, Rolfe PA, Heisler LE, Chin B, et al. 2010. Genotype to phenotype: a complex problem. Science 328:469.

Giaever G, Chu AM, Ni L, Connelly C, Riles L, Véronneau S, Dow S, Lucau-Danila A, Anderson K, André B, et al. 2002. Functional profiling of the Saccharomyces cerevisiae genome. Nature 418:387–391.

Glass JI, Assad-Garcia N, Alperovich N, Yooseph S, Lewis MR, Maruf M, Hutchison CA 3rd, Smith HO, Venter JC. 2006. Essential genes of a minimal bacterium. Proc. Natl. Acad. Sci. U. S. A. 103:425–430.

Harris MA, Rutherford KM, Hayles J, Lock A, Bähler J, Oliver SG, Mata J, Wood V. 2022. Fission stories: using PomBase to understand Schizosaccharomyces pombe biology. Genetics [Internet] 220. Available from: http://dx.doi.org/10.1093/genetics/iyab222

Hart GT, Lee I, Marcotte ER. 2007. A high-accuracy consensus map of yeast protein complexes reveals modular nature of gene essentiality. BMC Bioinformatics 8:236.

Hart T, Chandrashekhar M, Aregger M, Steinhart Z, Brown KR, MacLeod G, Mis M, Zimmermann M, Fradet-Turcotte A, Sun S, et al. 2015. High-Resolution CRISPR Screens Reveal Fitness Genes and Genotype-Specific Cancer Liabilities. Cell 163:1515–1526.

Hutchison CA 3rd, Chuang R-Y, Noskov VN, Assad-Garcia N, Deerinck TJ, Ellisman MH, Gill J, Kannan K, Karas BJ, Ma L, et al. 2016. Design and synthesis of a minimal bacterial genome. Science 351:aad6253.

Huttenhower C, Hibbs M, Myers C, Troyanskaya OG. 2006. A scalable method for integration and functional analysis of multiple microarray datasets. Bioinformatics 22:2890–2897.

Hwang Y-C, Lin C-C, Chang J-Y, Mori H, Juan H-F, Huang H-C. 2009. Predicting essential genes based on network and sequence analysis. Mol. Biosyst. 5:1672–1678.

Iorio F, Bernardo-Faura M, Gobbi A, Cokelaer T, Jurman G, Saez-Rodriguez J. 2016. Efficient randomization of biological networks while preserving functional characterization of individual nodes. BMC Bioinformatics 17:542.

Jordan DM, Frangakis SG, Golzio C, Cassa CA, Kurtzberg J, Task Force for Neonatal Genomics, Davis EE, Sunyaev SR, Katsanis N. 2015. Identification of cis-suppression of human disease mutations by comparative genomics. Nature 524:225–229.

Jordan IK, King Jordan I, Rogozin IB, Wolf YI, Koonin EV. 2002. Essential Genes Are More Evolutionarily Conserved Than Are Nonessential Genes in Bacteria. Genome Research [Internet] 12:962–968. Available from: http://dx.doi.org/10.1101/gr.87702

Juhas M, Eberl L, Glass JI. 2011. Essence of life: essential genes of minimal genomes. Trends Cell Biol. 21:562–568.

Kanehisa M, Sato Y, Kawashima M, Furumichi M, Tanabe M. 2016. KEGG as a reference resource for gene and protein annotation. Nucleic Acids Res. 44:D457–D462.

Kim D-U, Hayles J, Kim D, Wood V, Park H-O, Won M, Yoo H-S, Duhig T, Nam M, Palmer G, et al. 2010. Analysis of a genome-wide set of gene deletions in the fission yeast Schizosaccharomyces pombe. Nat. Biotechnol. 28:617–623.

van Leeuwen J, Pons C, Mellor JC, Yamaguchi TN, Friesen H, Koschwanez J, Ušaj MM, Pechlaner M, Takar M, Ušaj M, et al. 2016. Exploring genetic suppression interactions on a global scale. Science [Internet] 354. Available from: http://dx.doi.org/10.1126/science.aag0839

van Leeuwen J, Pons C, Tan G, Wang JZ, Hou J, Weile J, Gebbia M, Liang W, Shuteriqi E, Li Z, et al. 2020. Systematic analysis of bypass suppression of essential genes. Mol. Syst. Biol. 16:e9828.

Li G, Ji B, Nielsen J. 2019. The pan-genome of Saccharomyces cerevisiae. FEMS Yeast Res. [Internet] 19. Available from: http://dx.doi.org/10.1093/femsyr/foz064

Li J, Wang H-T, Wang W-T, Zhang X-R, Suo F, Ren J-Y, Bi Y, Xue Y-X, Hu W, Dong M-Q, et al. 2019. Systematic analysis reveals the prevalence and principles of bypassable gene essentiality. Nat. Commun. 10:1002.

Li L, Stoeckert CJ Jr, Roos DS. 2003. OrthoMCL: identification of ortholog groups for eukaryotic genomes. Genome Res. 13:2178–2189.

Liu G, Yong MYJ, Yurieva M, Srinivasan KG, Liu J, Lim JSY, Poidinger M, Wright GD, Zolezzi F, Choi H, et al. 2015. Gene Essentiality Is a Quantitative Property Linked to Cellular Evolvability. Cell 163:1388–1399.

Lloyd JP, Seddon AE, Moghe GD, Simenc MC, Shiu S-H. 2015. Characteristics of Plant Essential Genes Allow for within- and between-Species Prediction of Lethal Mutant Phenotypes. Plant Cell 27:2133–2147.

Luo H, Gao F, Lin Y. 2015. Evolutionary conservation analysis between the essential and nonessential genes in bacterial genomes. Sci. Rep. 5:13210.

Mackay TFC. 2014. Epistasis and quantitative traits: using model organisms to study gene–gene interactions. Nature Reviews Genetics [Internet] 15:22–33. Available from: http://dx.doi.org/10.1038/nrg3627

Meldal BHM, Pons C, Perfetto L, Del-Toro N, Wong E, Aloy P, Hermjakob H, Orchard S, Porras P. 2021. Analysing the yeast complexome-the Complex Portal rising to the challenge. Nucleic Acids Res. 49:3156–3167.

Meyers RM, Bryan JG, McFarland JM, Weir BA, Sizemore AE, Xu H, Dharia NV, Montgomery PG, Cowley GS, Pantel S, et al. 2017. Computational correction of copy number effect improves specificity of CRISPR-Cas9 essentiality screens in cancer cells. Nat. Genet. 49:1779–1784.

Mi H, Ebert D, Muruganujan A, Mills C, Albou L-P, Mushayamaha T, Thomas PD. 2021. PANTHER version 16: a revised family classification, tree-based classification tool, enhancer regions and extensive API. Nucleic Acids Res. 49:D394–D403.

Myers CL, Barrett DR, Hibbs MA, Huttenhower C, Troyanskaya OG. 2006. Finding function: evaluation methods for functional genomic data. BMC Genomics 7:187.

Narasimhan K, Lambert SA, Yang AWH, Riddell J, Mnaimneh S, Zheng H, Albu M, Najafabadi HS, Reece-Hoyes JS, Fuxman Bass JI, et al. 2015. Mapping and analysis of Caenorhabditis elegans transcription factor sequence specificities. Elife [Internet] 4. Available from: http://dx.doi.org/10.7554/eLife.06967

Ng PC, Henikoff S. 2001. Predicting deleterious amino acid substitutions. Genome Res. 11:863–874.

Peter J, De Chiara M, Friedrich A, Yue J-X, Pflieger D, Bergström A, Sigwalt A, Barre B, Freel K, Llored A, et al. 2018. Genome evolution across 1,011 Saccharomyces cerevisiae isolates. Nature 556:339–344.

Rancati G, Moffat J, Typas A, Pavelka N. 2018. Emerging and evolving concepts in gene essentiality. Nat. Rev. Genet. 19:34–49.

Roemer T, Jiang B, Davison J, Ketela T, Veillette K, Breton A, Tandia F, Linteau A, Sillaots S, Marta C, et al. 2003. Large-scale essential gene identification in Candida albicans and applications to antifungal drug discovery. Mol. Microbiol. 50:167–181.

Sanchez MR, Payen C, Cheong F, Hovde BT, Bissonnette S, Arkin AP, Skerker JM, Brem RB, Caudy AA, Dunham MJ. 2019. Transposon insertional mutagenesis in reveals -acting effects influencing species-dependent essential genes. Genome Res. 29:396–406.

Sardiu ME, Gilmore JM, Carrozza MJ, Li B, Workman JL, Florens L, Washburn MP. 2009. Determining protein complex connectivity using a probabilistic deletion network derived from quantitative proteomics. PLoS One 4:e7310.

Segal ES, Gritsenko V, Levitan A, Yadav B, Dror N, Steenwyk JL, Silberberg Y, Mielich K, Rokas A, Gow NAR, et al. 2018. Gene Essentiality Analyzed by Transposon Mutagenesis and Machine Learning in a Stable Haploid Isolate of. MBio [Internet] 9. Available from: http://dx.doi.org/10.1128/mBio.02048-18

Sonnhammer ELL, Östlund G. 2015. InParanoid 8: orthology analysis between 273 proteomes, mostly eukaryotic. Nucleic Acids Res. 43:D234–D239.

Takeda A, Saitoh S, Ohkura H, Sawin KE, Goshima G. 2019. Identification of 15 New Bypassable Essential Genes of Fission Yeast. Cell Struct. Funct. 44:113–119.

UniProt Consortium. 2021. UniProt: the universal protein knowledgebase in 2021. Nucleic Acids Res. 49:D480–D489.

Wang T, Birsoy K, Hughes NW, Krupczak KM, Post Y, Wei JJ, Lander ES, Sabatini DM. 2015. Identification and characterization of essential genes in the human genome. Science 350:1096–1101.

Wei W-H, Hemani G, Haley CS. 2014. Detecting epistasis in human complex traits. Nature Reviews Genetics [Internet] 15:722–733. Available from: http://dx.doi.org/10.1038/nrg3747

Woodford N, Ellington MJ. 2007. The emergence of antibiotic resistance by mutation. Clin. Microbiol. Infect. 13:5–18.

Wood V, Harris MA, McDowall MD, Rutherford K, Vaughan BW, Staines DM, Aslett M, Lock A, Bähler J, Kersey PJ, et al. 2012. PomBase: a comprehensive online resource for fission yeast. Nucleic Acids Res. 40:D695–D699.

Zhang X, Acencio ML, Lemke N. 2016. Predicting Essential Genes and Proteins Based on Machine Learning and Network Topological Features: A Comprehensive Review. Front. Physiol. 7:75.

